# Ubiquitous Forbidden Order in R-group classified protein sequence of SARS-CoV-2 and other viruses

**DOI:** 10.1101/2020.08.21.261289

**Authors:** Pratibha, C. Shaju, Kamal

**Affiliations:** Department of Mathematics, Indian Institute of Technology Roorkee, India; Department of Earth Sciences, Indian Institute of Technology Roorkee, India; Department of Earth Sciences, Centre of Excellence in Disaster Mitigation and Management, Indian Institute of Technology Roorkee, India

**Keywords:** iterated function systems, amino acids, genome, protein sequence, side-chain, chaos game, forbidden order

## Abstract

Each amino acid in a polypeptide chain has a distinctive R-group associated with it. We report here a novel method of species characterization based upon the order of these R-group classified amino acids in the linear sequence of the side chains associated with the codon triplets. In an otherwise pseudo-random sequence, we search for forbidden combinations of *k*th order. We applied this method to analyze the available protein sequences of various viruses including SARS-CoV-2. We found that these ubiquitous forbidden orders (UFO) are unique to each of the viruses we analyzed. This unique structure of the viruses may provide an insight into viruses’ chemical behavior and the folding patterns of the proteins. This finding may have a broad significance for the analysis of coding sequences of species in general.

## Main

Each trinucleotide or a codon triplet in a genome sequence represents an amino acid which are the basic building blocks of proteins. The amino acids have an amine group at one end and a distinctive side chain on the other. Although the backbone of all amino acids is the same, the side chains are not. The order, in which the amino acids are aligned, is governed by the genetic code.

One of the fundamental tasks of genomics is to compare these coding sequences for phylogenetic analysis. The comparison may be carried out either by sequential alignment [1,2] or through an alignment-free approach [3, 4, 5].One of the alignment-free approaches is based on a driven iterated function system (IFS) called chaos game representation (CGR) which has become popular due to its compact representation of the whole genome sequence in the graphical form [6,7,8,9,10]. Recently, a modified form of this approach has been used to study the SARS-CoV-2 sequences [11, 12]. Attempts were also made to use CGR method to analyze the protein sequences directly [13] or through their side chains [14].

All the available studies in genomic data using driven IFS focus on the order of either the nucleotides (in case of DNA/RNA) or the amino acids (in case of proteins). We decided to look at the other side of the coin and propose to study the forbidden combinations in a sequence of amino acids.

An amino acid in a polypeptide chain has a distinctive R-group associated with it. We characterize human coronaviruses based upon the order in the linear sequence of the side chains associated with codon triplets. In an otherwise pseudo-random sequence, we search for forbidden combinations of *k*^th^ order. The results indicate what nature has decided not to do rather than what to do. We found that these forbidden orders are ubiquitous to each of the viruses we analyzed. These ubiquitous forbidden orders (UFO) are unique structures of the viruses that may provide an insight into viruses’ chemical behavior and the folding patterns of the proteins. This finding may have a broad significance for the analysis of coding sequences of species in general.

## Material and Methods

Coding sequences of species were downloaded from the National Centre for Bioinformatics (NCBI) website (https://www.ncbi.nlm.nih.gov/). The species used in the study, with their accession id, are SARS-CoV-2 (NC_045512.2), SARS-COV-1 (NC_004718.3), MERS (NC_019843.3), HKU1 (NC_006213.1), NL63 (NC_005831.2), OC43 (NC_006213.1), 229E (NC_002645.1), Bat coronavirus RaTG13 isolate (MN996532.1), Marburg (NC_024781.1), Rabies (NC_001542.1), Variola (smallpox) (NC_001611.1), Influenza H1N1 (NC_026431 – 38.1, 8 segments), Hantavirus (NC_005215 – 17.1, 3 segments), Human Rota virus (NC_021541 – 51.1, 11 segments), Bovine corona (NC_003045.1), Betacorona England1 (NC_038294.1), Nipah Henipavirus (NC_002728.1), Hendra Henipavirus (NC_001906.3), Measles morbillivirus (NC_001498.1), Zika (NC_035889.1), Polio (NC_9002058.3), Hepatitis A (NC_001489.1), Mumps (NC_002200.1), Human parainfluenza (NC_021928.1), HIV Congo 1987 (MH705134.1), and Human Adenovirus (NC_012959.1).

The codon triplets of four bases (A, G, C, and T) in the sequence can have 64 possible combinations that form 20 amino acids. Each of these amino acids is associated with a side chain that controls the folding patterns of the protein and its chemical behavior. Based on the chemical properties of the side chain, each codon triplet can be sub-classified as Non-polar (N), uncharged polar (P), acidic (A), or basic (B). A MATLAB code reads the sequence and classifies each amino acid triplet as its respective R-group side chain. Based on the chemical properties of the side chain of an amino acid in a coding sequence of a species, we generated a sequence of N (non-polar), P (polar), A (acidic), and B (basic). A new sequence of four symbols, N, B, A, and P is thus created from the protein sequence of the species that looks something like this - NBNNPAAABBNPNNNPABBABABAA…………… where each letter represents amino acid with one of the chemical properties listed above. This R-group sequence is used to obtain CGR plots of any given protein sequence of a species (Figure 1). Theoretically, all combinations of N, B, A, and P are possible in this sequence. In this study, we look at the sequence from a different perspective. Instead of studying what is present in the protein sequence, we decided to analyze what is absent from the sequence.

**Figure 1.**
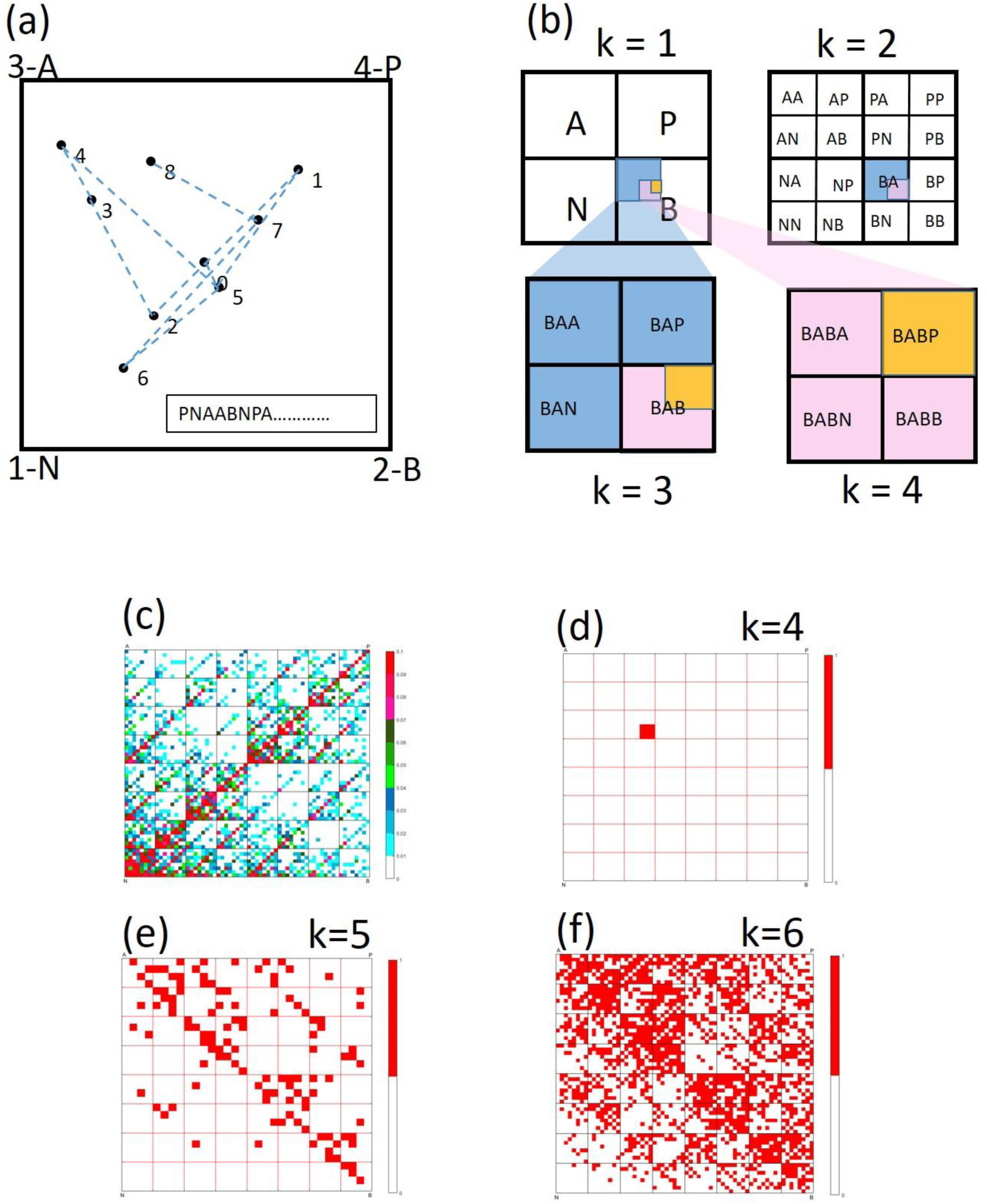
a. The actual path of the IFS as governed by the corresponding side-chain of the amino acid in a coding sequence. The vertices of the square correspond to the polarity of the side chain, Non-polar (N), Basic (B), Acidic (A), and uncharged Polar (P). We begin at the center of the square and move halfway towards a vertex depending on the letter occurrence in the R-group sequence. We repeat this for each base as they appear in the R-group sequence. Finally, we get a plot of points according to the order of the R-group member in the sequence. These points will occupy one of the addresses in the square as depicted in Figure 1b. b. The address map of the unit square NBAP (defined in a). Longer addresses denote smaller subsquares. The empty subsquares in a driven IFS indicate a forbidden combination of that order. c. Percentage CGR plot (PC-plot) of Human Adenovirus for k = 6. The colors indicate the intensity of points in a subsquare of order 6. d. e. f. Ubiquitous Forbidden Order (UFO) plots of Human Adenovirus for combinations of order (k) 4, 5, and 6. The vertices in the squares are the same as depicted in figure 1a The red color indicates that the corresponding address is forbidden. For example, In figure 1d, the address ABAB is forbidden.

### CGR of a driven IFS

A protein sequence *X(k)* can be considered as a string composed of N, P, A, and B.

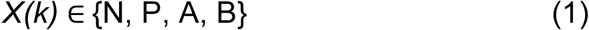

We consider a unit square *U* and name corners *C*_*i*_ (*i*=1,2,3,4) as N, B, A, and P respectively, which corresponds to the value of *X(k)*. The initial point P_(0)_ is the midpoint of the square. Now the second point P_(1)_ is the midpoint between P_(0)_ and C_X(1)_. In General, P_(k)_ is plotted as the midpoint between P_(k-1)_ and C_X(k)_ [6].

After plotting the genetic sequence *X* in unit square *U*, the unit square is divided into 2^k^ x 2^k^ sub squares; each sub-square represents a unique sub-sequence of length k. An example of the movement of points in CGR is shown with the first eight members of the data sequence (PNAABNPA….) in Figure 1a. An example of addresses of the sub-squares for different orders (k =, 1, 2, 3, and 4) is given in Figure 1b.

### PC-plots

To make these plots, the percentage of points plotted in sub-square is calculated. This percentage value represents the intensity of points in each sub-square. After plotting points by CGR and dividing the unit square into 2^k^ x 2^k^ sub squares, each sub-square is color-filled based on the calculated intensity values. Figure 1c shows the percentage-CGR plot made for the Human Adenovirus for k=6. The existing literature presents studies of these plots in a phylogenetic analysis of species [6,7,9,10,12]. We take a step further but in the opposite direction and look for those combinations of N, B, A, and P, which are ubiquitously forbidden by nature for a given length (k).

### UFO plots

We generated driven iterated function system PC-plots for the species and look for the forbidden combinations. The UFO plots are complimentary to the PC-plots. Here we look at the sub-sequences of a particular length (order) of N, B, A, and P which are prohibited by the species. The Forbidden address in a UFO plot and the actual order that is forbidden by nature are counter-intuitive and are opposite to each other.For example, if ABAB does not occur in the protein sequence of Human Adenovirus as shown in Figure 1d, then we say that the 4^th^ order combination BABA is forbidden, i.e., Basic, Acidic, Basic and Acidic side chains, in that order, are not possible in the protein sequence. This also means that if at a point in human adenovirus, the side chains are basic, acidic, and basic at the 3^rd^ order, then the next side chain can only be polar, non-polar, or basic. It cannot be acidic.

The forbidden order in the plots can only be visualized for lower orders. It can be seen that Figures 1e and 1f are becoming more and more chaotic as the value of k increases and are difficult to analyze.

## Results

Next, we analyzed protein sequences of 26 viruses (Figures 2, 3, and Supplementary Figures 1 – 4) to search for a ubiquitous forbidden order in each one of them. Our purpose was to find some clues for uniquely analyzing the SARS-CoV-2 to handle the COVID-19 pandemic.

**Figure 2.**
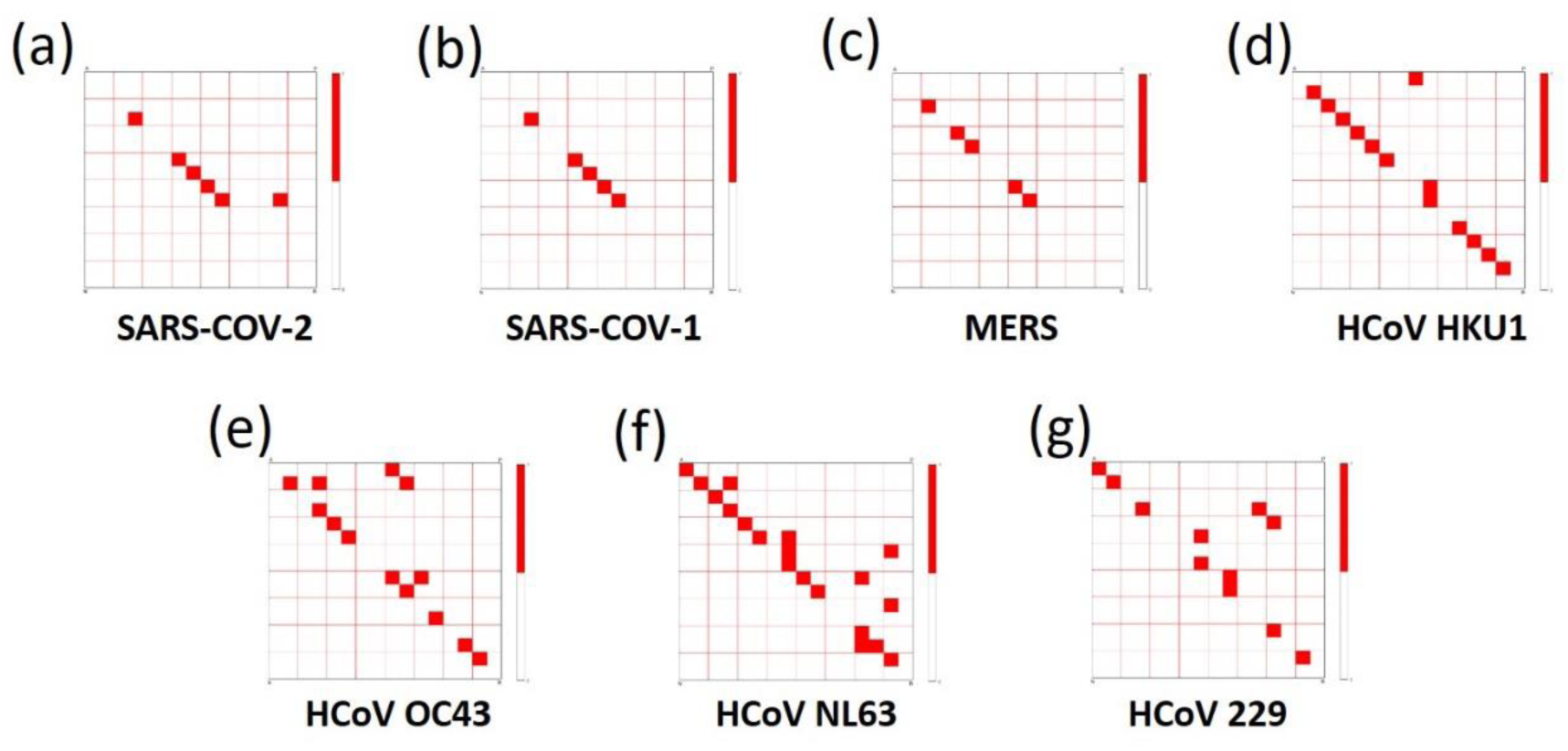
A comparison of the 4^th^ level ubiquitous forbidden order (UFO) of the human coronaviruses. Here all human coronavirus UFOs are shown in the order they were discovered starting from the present, namely, SARS-CoV-2 (a), SARS-CoV-1 (b), MERS (c), HCoV HKU1 (d), HCoV NL63 (e), HCov OC43 (f), and HCoV 229E (g). Although the UFO plot is unique for each virus, they seem to optimize their evolution in time.

**Figure 3.**
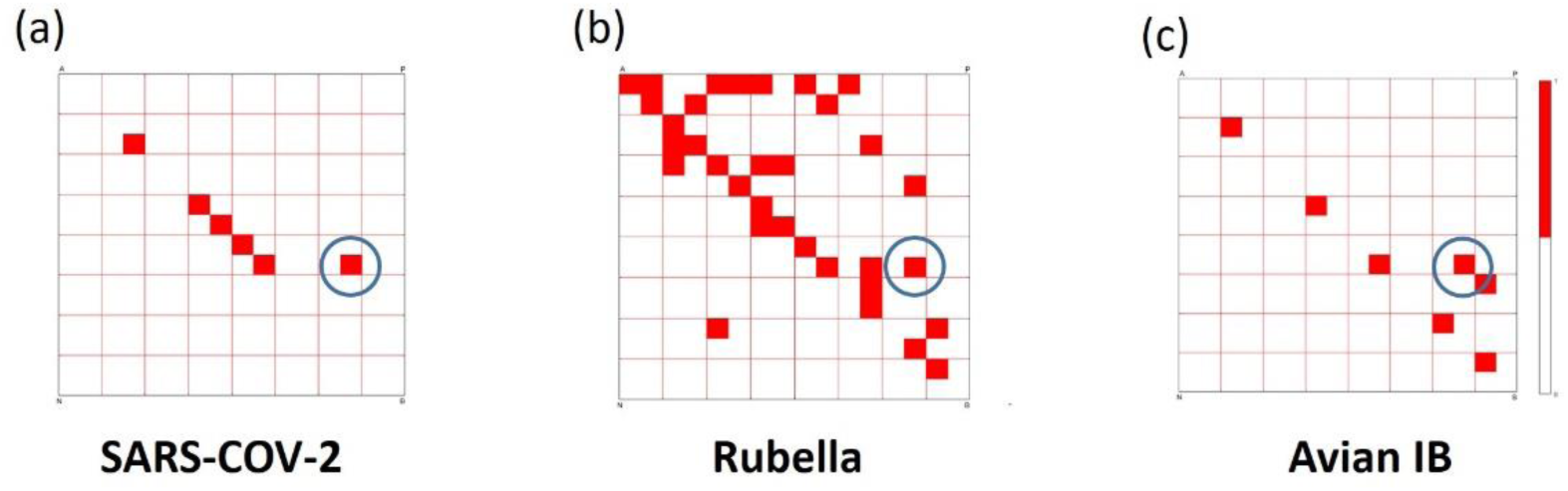
A comparison of the ubiquitous forbidden order (UFO) of the species. Here three viruses’ UFO plots are shown, namely, SARS-CoV-2 (a), Rubella (b), and Avian IB (c). Although the UFO portrait is unique for each virus, a common feature to these three is the address BPAB (encircled in blue), which corresponds to the forbidden order BAPB in the amino acid sequence.

Figure 2 shows the 4^th^ level UFO plots of seven coronaviruses infecting humans. From UFO plots, the viruses seem to be getting optimized with time. The earlier coronaviruses, 229E, OC43, NL63, and HKU1 (Figure 1 d, e, f, and g), which are also the mild coronaviruses, have a lot of forbidden addresses in the amino acid polypeptide chain. With evolution, the structure seems to be getting simpler for MERS, SARS-CoV-1, and SARS-CoV-2(Figure 1 a, b, and c). It appears that nature prefers to be simple and less complex to be able to survive and evolve.

A close examination of UFO plots of SARS-CoV-2 (Figure 2a) and SARS-CoV-1 (Figure 2b) reveals that the evolution of the SARS-CoV-2 from SARS-CoV-1 is quite straight forward. One just has to prohibit a particular order BAPB in the SARS-CoV-1 protein sequence to get a SARS-CoV-2 strain. This opens up a point of discussion whether this formation is possible in a laboratory setting or not. As non-biologists, we cannot comment on their origin, though our results provide an alternative approach for further exploration by the subject specialists.

Among the 26 viruses studied, we noted that the forbidden order BPAB is unique to SARS CoV-2, Rubella, and Avian IB (Figure 3). A forbidden order BPAB in the UFO plot means that the 4^th^ order combination BAPB is prohibited by nature, i.e., a basic side chain(B), followed by an acidic side chain(A), followed by an uncharged polar side chain(P) cannot be followed by a basic side chain(B). This rule is found to be followed only by the SARS-CoV-2, Rubella, and the Avian Infectious Bronchitis (AIB) sequences among all viruses studied by us. The whole forbidden structure of SARS-CoV-2 is an inherent part of the Rubella plot and partly of the AIB plot.

This feature in the SARS-CoV-2 protein sequence may be a pointer to support the idea of using the MMR vaccine in COVID-19 as floated by Fidel and Noverr [15] and the use of recombinant ACE2 by Kruse[16] as a preventive measure to reduce the inflammation in COVID-19 patients.The use of drugs/vaccines for existing viruses in managing untreatable viral diseases has been suggested by others also [17, 18] including for COVID-19 [19].

Next, we tried to analyze the studies on the evolutionary origin of the SARS-CoV-2 and its comparison to the Bat coronavirus RaTG13 isolate [20,21]. We found that at R-group classified sequences of N, B, A, and P in these two samples are identical up to level 4 of the amino acid ordering in the protein structures (Figures 4a, d, g). However, at the next level, i.e. k=5, the differences between the two protein sequences start to emerge (Figures 4b, e, h) and become quite clear at the 6^th^ level of ordering (Figures 4c, f, i). This supports the findings of Wrobel *et.al*.[21].

**Figure 4.**
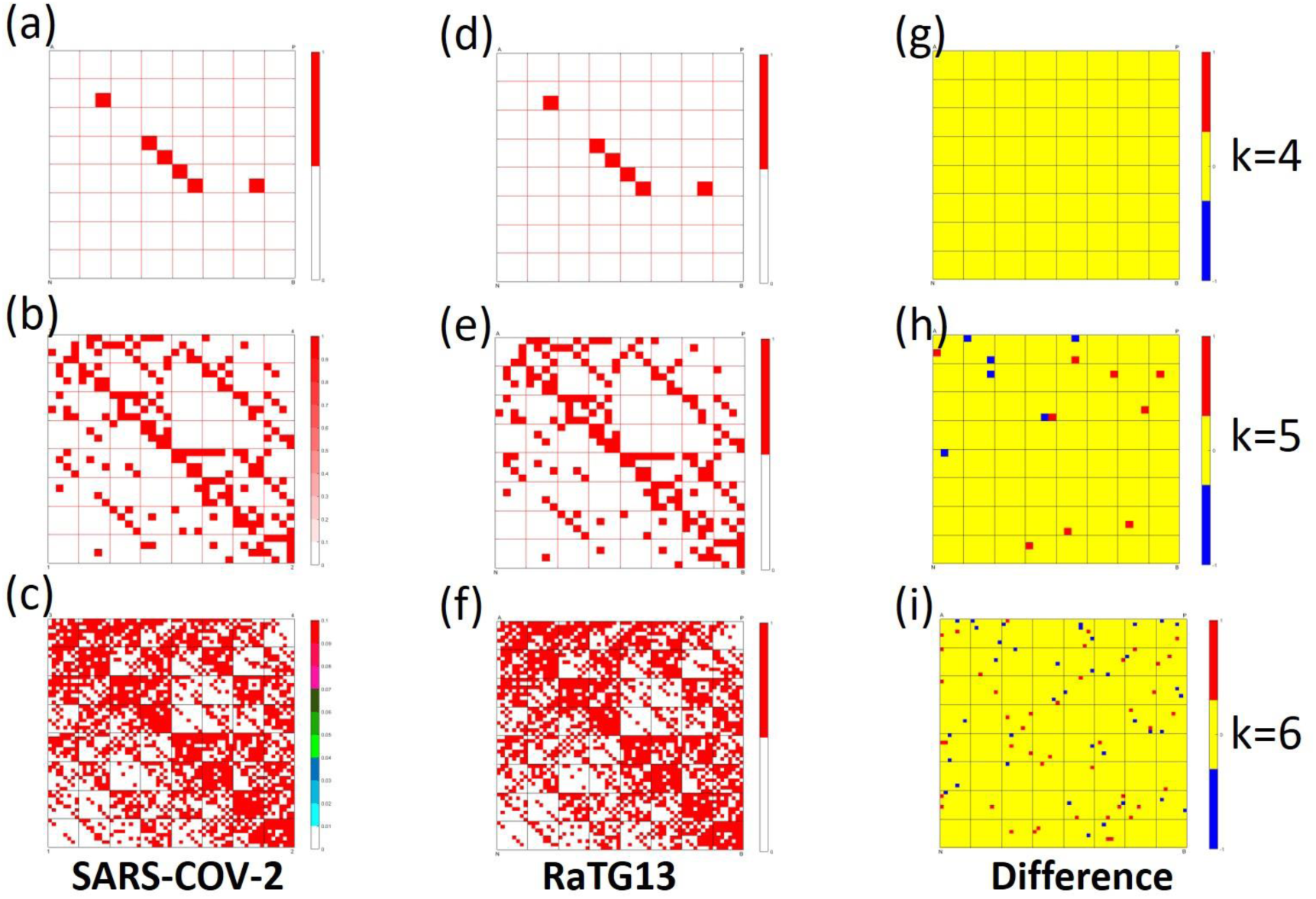
A comparison of UFO plots of SARS-CoV-2 and Bat RaTG13 coronavirus isolate. a. b. c. are UFO of SARS-CoV-2 and d. e. f. are UFO for RaTG13 isolate for order k = 4. 5, and 6 respectively. g. h. i. are the difference plots of these for the corresponding order. The difference plots denote the key addresses which are forbidden in the first plot but present in the second with red filled sub-squares whereas the blue sub-squares indicate the addresses that are forbidden in the second plot but are allowed in the first one. Note the increase in complexity with the increase in order. There is no evident difference between the SARS-CoV-2 and its closest relative[20] RaTG13 structures at the 4^th^ order (Figure 4g), but the higher-order comparison (Figures 4h, i) reveals key differences between these two viruses.

## Conclusions

Recently, Flies *et.al*.[22] emphasized shifting the focus of immunological research to new models and interdisciplinary studies. We have looked at the amino acid sequences, as non-biologists, from a different angle based on the chemical properties of the side chain of an amino acid in a coding sequence of a species. The forbidden order BAPB is unique to SARS CoV-2. A basic side chain, followed by an acidic side chain, followed by an uncharged polar side chain can not be followed by a basic side chain. This rule is found to be followed only by the SARS-CoV-2, Rubella, and the Avian Infectious Bronchitis among the viruses studied by us.

This study of the forbidden order of R-group side chains in a protein sequence opens up new directions for microbiologists to study coding sequences.The consequences of the forbidden order to the properties of a protein are yet to be ascertained.

## References

1. Zhi Qi, Sy Redding, Ja Yil Lee, Bryan Gibb, YoungHo Kwon, Hengyao Niu, William A. Gaines, Patrick Sung, Eric C. Greene, DNA Sequence Alignment by Microhomology Sampling during Homologous Recombination, Cell 160, 856–869, https://doi.org/10.1016/j.cell.2015.01.029 (2015).

2. Altschul, S. F. et al. Gapped blast and psi-blast: a new generation of protein database search programs. Nucleic Acids Research 25, 3389–402 (1997).

3. Deng, M., Yu, C., Liang, Q., He, R. L. & Yau, S. S.-T. A novel method of characterizing genetic sequences: genome space with biological distance and applications. PLoS ONE 6, e17293 (2011).

4. Hoang, T., Yin, C. & Yau, S. S.-T. Numerical encoding of dna sequences by chaos game representation with application in similarity comparison. Genomics 108, 134–142 (2016).

5. Rohling S, Linne A, Schellhorn J, Hosseini M, Dencker T, Morgenstern B The number of k-mer matches between two DNA sequences as a function of k and applications to estimate phylogenetic distances. PLoS ONE 15(2): e0228070. https://doi.org/10.1371/journal.pone.0228070 (2020).

6. Jeffrey, H. J. Chaos game representation of gene structure. Nucleic Acids Research 18, 2163–2170 (1990).

7. Deschavanne PJ1, Giron A, Vilain J, Fagot G, Fertil B. Genomic signature: characterization and classification of species assessed by chaos game representation of sequences. Mol Biol Evol. 16(10).,1391–1399, (1999). doi: 10.1093/oxfordjournals.molbev.a02604

8. Almeida, J. S., Carriço, J. A., Maretzek, A., Noble, P. A. & Fletcher, M. Analysis of genomic sequences by Chaos Game Representation. Bioinformatics 17, 429–437 (2001).

9. C. Stan, C.P. Cristescu, E.I. Scarlat, Similarity analysis for DNA sequences based on chaos game representation case study: The albumin, J. Theoret. Biol., 267, 513–518, (2010)

10. Lichtblau, D. Alignment-free genomic sequence comparison using FCGR and signal processing. BMC Bioinformatics 20, 742 https://doi.org/10.1186/s12859-019-3330-3, (2019).

11. Pratibha, Shaju C., Gupta. A., Kamal. (2020). A categorization of COVID-19 treatment strategies: A modified chaos game representation (CGR) analysis of genome sequences for thirty-two pathogens. COVID-19 virtual conference, AIDS 2020, an IAS virtual conference, SanFrancisco, USA. July 10-11, 2020.

12. Randhawa GS, Soltysiak MPM, El Roz H, de Souza CPE, Hill KA, Kari L (2020). Machine learning using intrinsic genomic signatures for rapid classification of novel pathogens: COVID-19 case study. PLoS ONE 15(4): e0232391. https://doi.org/10.1371/journal.pone.0232391

13. Zeju Sun, Shaojun Pei, Rong Lucy He, Stephen S.-T. Yau, 2020. A novel numerical representation for proteins: Three-dimensional Chaos Game Representation and its Extended Natural Vector, Computational and Structural Biotechnology Journal, 18, 1904–1913, https://doi.org/10.1016/j.csbj.2020.07.004

14. Yu, Z.G., Anh, V., Lau, K.S., 2004. Chaos game representation of protein sequences based on the detailed HP model and their multifractal and correlation analyses. J. Theor. Biol. 226, 341–348

15. Fidel PL, Jr., Noverr MC. 2020. Could an unrelated live attenuated vaccine serve as a preventive measure to dampen septic inflammation associated with COVID-19 infection? mBio 11:e00907–20. https://doi.org/10.1128/mBio.00907-20.

16. Kruse RL. 2020. Therapeutic strategies in an outbreak scenario to treat the novel coronavirus originating in Wuhan, China. F1000Res. 9:72. Published 2020 Jan 31. https://doi.org/10.12688/f1000research.22211.2

17. Aaby P, Benn CS. 2019. Developing the concept of beneficial nonspecific effect of live vaccines with epidemiological studies. Clin. Microbiol Infect 25:1459 –1467. https://doi.org/10.1016/j.cmi.2019.08.011.

18. Moorlag S, Arts RJW, van Crevel R, Netea MG. 2019. Non-specific effects of BCG vaccine on viral infections. Clin Microbiol Infect 25:1473–1478. https://doi.org/10.1016/j.cmi.2019.04.020

19. Gordon, D.E., Jang, G.M., Bouhaddou, M. et al. A SARS-CoV-2 protein interaction map reveals targets for drug repurposing. Nature 583, 459–468 (2020). https://doi.org/10.1038/s41586-020-2286-9

20. Boni, M.F., Lemey, P., Jiang, X. et al. 2020. Evolutionary origins of the SARS-CoV-2 sarbecovirus lineage responsible for the COVID-19 pandemic. Nature Microbiol. https://doi.org/10.1038/s41564-020-0771-4

21. Wrobel, A.G., Benton, D.J., Xu, P. et al. 2020. SARS-CoV-2 and bat RaTG13 spike glycoprotein structures inform on virus evolution and furin-cleavage effects. Nat Struct Mol Biol 27, 763–767. https://doi.org/10.1038/s41594-020-0468-7

22. Flies, A.S. 2020. Rewilding immunology, Science, 369 (6499), 37–38. https://doi.org/10.1126/science.abb8664

